# The Importance of Mentors and How to Handle More Than One Mentor

**DOI:** 10.1101/2021.11.29.469764

**Authors:** Andrea G. Marshall, Lillian J. Brady, Caroline B. Palavicino-Maggio, Kit Neirkirk, Zer Vue, Heather Beasley, Edgar Garza-Lopez, Sandra Murray, Denise Martinez, Haysetta Shuler, Elsie C. Spencer, Derrick Morton, Antentor Hinton

**Author notes:** denotes co-first author (Andrea G. Marshall, Lillian J. Brady). denotes co-senior author (Haysetta Shuler, Elsie C. Spencer, Derrick Morton, Antentor Hinton).

## Abstract

**Introduction:** Working with multiple mentors is a critical way for students to expand their network, gain opportunities, and better prepare for future scholastic or professional ventures. However, students from underrepresented groups (UR) are less likely to be mentored or have access to mentors, particularly in science, technology, engineering, and mathematics (STEM) fields. We developed and implemented a workshop, to provide the necessary foundation for students to be better prepared for establishing future mentorships throughout graduate and professional school.

**Methods:** Faculty well-versed in the area of effective mentorship from multiple universities developed and delivered a 1.5-hour workshop to address the roles of a mentor, especially when it comes to UR students, and how students may effectively work with multiple mentors. This workshop was delivered to a group of students from the HBCU Winston Salem State University, and a pre/post-Likert scale-based survey was administered.

**Results:** We analyzed the raw data with nonparametric tests for comparison within paired samples. Wilcoxon matched-pairs and signed-rank tests showed statistically significant growth in student self-ratings related to the workshop learning objectives.

**Conclusions:** The “How to Handle More than One Mentor to Achieve Excellence” workshop was well received as a component of pre-graduate and pre-professional training. Incorporating workshops like this may increase student preparedness around developing and cultivating healthy mentorship relationships throughout STEM training.

**EDUCATIONAL OBJECTIVES:** By the end of this workshop, learners will be able to:

1. Describe the role of mentors in developing the next generation of trainees.
2. Describe current research on mentorship among underrepresented populations.
3. Apply skills on effective communication needed in the development of successful mentorship relationships.
4. Work with multiple mentors at one time while maintaining solid professional and personal relationships.

## INTRODUCTION

Mentors are crucial for supporting students of all levels, be they undergraduate, post-undergraduate, or beyond. Making use of a mentor and working with multiple mentors is a keyway for students to expand their network, gain opportunities, and better prepare for future scholastic or professional ventures. Studies have shown that effective mentorship can increase productivity by reducing stress and bettering mental health (Hund *et al*. 2018). Mentoring is an important step in increasing diversity, inclusion, and equality in science, technology, engineering, and mathematics (STEM) by increasing the involvement of students who may otherwise not consider STEM fields (National Academies of Sciences 2020). The importance of this is underscored by the fact that diverse individuals have been found to produce more novel findings that are relevant to a wider-audience (Ding *et al*. 2021). Ultimately, many underrepresented (UR) minorities may need assistance in their informal job skills, including networking and career development skills, and mentors can be important in bridging that gap by providing important personal and professional development opportunities throughout the mentee’s career (Thomas, Willis and Davis 2007). Unfortunately, studies have found that students from backgrounds traditionally UR in STEM lack exposure to or have less access to mentoring (Kalbfleisch and Davies 1991; Ragins 1997; Thomas, Willis and Davis 2007; McDonald and Westphal 2013). Therefore, we developed and implemented a workshop that aimed to address issues related to students making use of multiple mentors, gaining networking experience and opportunities. Thoroughly understanding the mentoring relationship cycle **(Figure 1)** as well as the importance of cultivating a healthy mentee-to-mentor relationship comes with its own challenges for both mentees and mentors. While some students may have an idea of what a mentor should be, they often lack the deeper understanding of what a mentor specifically is, what a mentor does, how to foster a healthy mentee-to-mentor relationship, and how to work with multiple mentors at one time (Saito and Blyth 1992; Amaral and Vala 2009). This workshop sought to better fill these gaps in knowledge to, ultimately, create students well equipped for healthy and productive mentorships.

**Figure 1.**
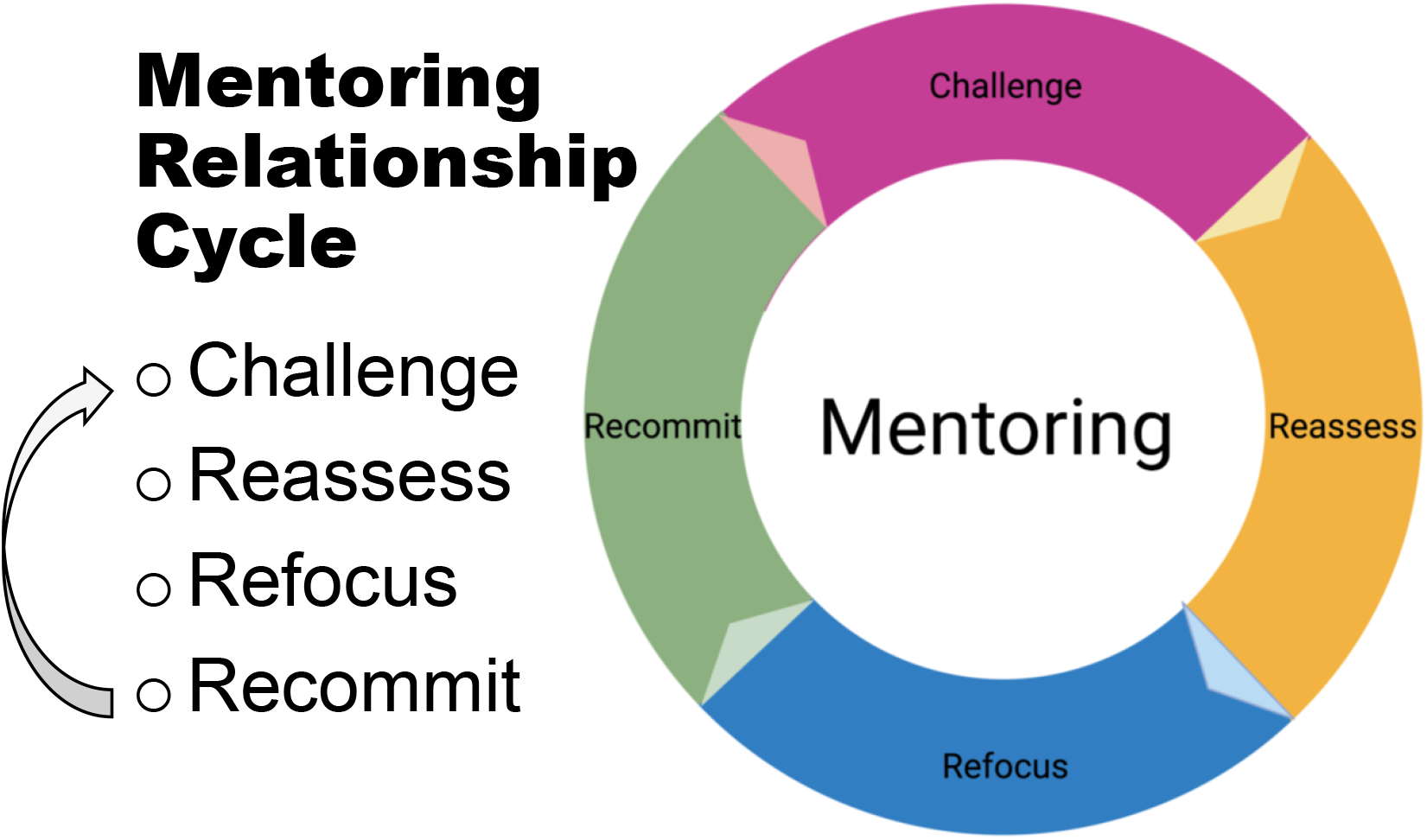
Graphic adapted from the workshop depicting the mentoring relationship cycle. Often in mentoring relationships, a <u>challenge</u> occurs that both mentee and mentor must work through to first <u>reassess</u> the mentee-to-mentor relationship, followed by a period of <u>refocusing</u> on the overall goals of the mentee-to-mentor relationship, and ending with an effort on the part of both mentee and mentor to <u>recommit</u> to the mentoring relationship before another challenge may arise.

The word mentor originated as the advisor of Telemachus in Homer’s odyssey (Sambunjak and Marušić 2009). Currently, the term ‘mentor’ can encompass a wide array of roles and actions. For example, one mentor may provide help through personal struggles but not professional struggles, while another may aid in training your technical skills but not with setting your career goals. In some cases, mentor is simply a title that describes the person for whom you work. Specifically, we define an effective mentor as one who is taking proper actions and making well-informed decisions to enact positive changes for both themselves and their mentee (Shuler *et al*. 2021). Therefore, it is critical for students to understand what constitutes an effective mentor and mentoring relationship.

A mentor can play multiple roles that span different scopes—it is not a one-size-fits-all position. A mentor can be a(n) coach, advisor, sponsor, and/or listener **(Figure 2)**. While each mentor typically has a defined role in their mentee’s life, the mentoring relationship is not static. For example, some mentorships will only exist within the context of the workplace, while others may develop into friendships or lifelong mentorships. As such, mentoring relationships may have variable lengths and exist for different, or multiple, purposes (Arnold and Johnson 1997; Rhodes *et al*. 2017). In general, however, a mentor is defined one who is a willing and trusted advisor who provides support, including emotional, academic, or a combination of the two (Hall *et al*. 2008; Haggard *et al*. 2011; Israel *et al*. 2014; Shuler *et al*. 2021). Importantly, a mentor can be anyone; however, that does not mean that a particular mentor is a good fit for just all mentees. The mentee and mentor must have a relationship that allows them to both work together effectively. This workshop defines what a mentor is and is not to help students better understand the broad scope of mentoring, with the goal of increasing effective mentoring relationships for UR students in STEM.

**Figure 2.**
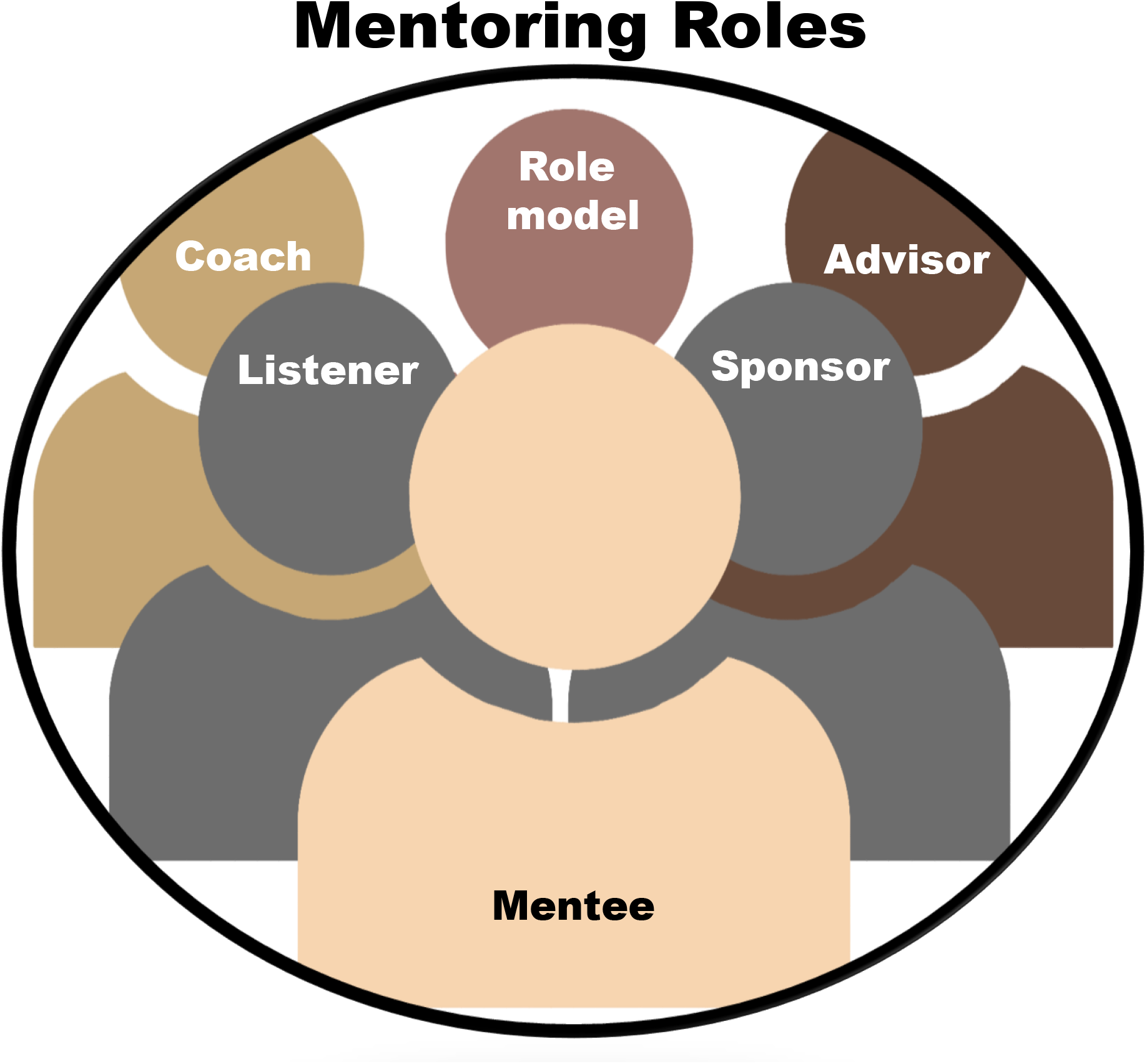
Infographic adapted from the workshop depicting the different roles of a mentor for underrepresented mentees which include but are not limited to being a(n) coach, advisor, sponsor, listener, and/or role model.

This seminar is also relevant for established scientists and educators, as it focuses on how mentors can maximize their effectiveness. Mentors should be aware of the various factors that affect success of UR students, such as stress, anxiety, available opportunities, and biases **(Figure 3)** (Shuler *et al*. 2021). As a result of this, mentors should work to share personal experiences with their mentees to foster an open and welcoming environment help build a bond with their mentee, while actively improving their ability to help their mentees overcome their unique struggles (Shuler *et al*. 2021). This workshop discussed these considerations along with offering solutions to mentors on how to deal with this. For UR mentees, they typically need to deal with stereotype threat – which may create favorable or unfavorable generalizations about a racial or gender group **(Figure 4)** (Hinton Jr *et al*. 2020). While there are many negative prejudices that UR must overcome to have a successful career in science, they must also combat the effects of people unwittingly causing trouble by considering students as ‘model minorities’. Categorizing as a “model minority” is the act of considering a certain minority class or individual as being based on a stereotype **(Figure 3)**. While this is both harmfully stereotyping other minorities, for those being classified as a model, it can create a pressure on the mentee to succeed (Wong and Halgin 2006; Kiang *et al*. 2017). Furthermore, even as these mentees progress, their struggles may be trivialized through tokenism, which is when institutions or groups may assume that an UR individual can serve as a voice for that entire population. As such, this workshop was multifaceted in both highlighting how to better mentor students while explaining the importance of mentors for potential mentees **(Figure 2)**.

**Figure 3.**
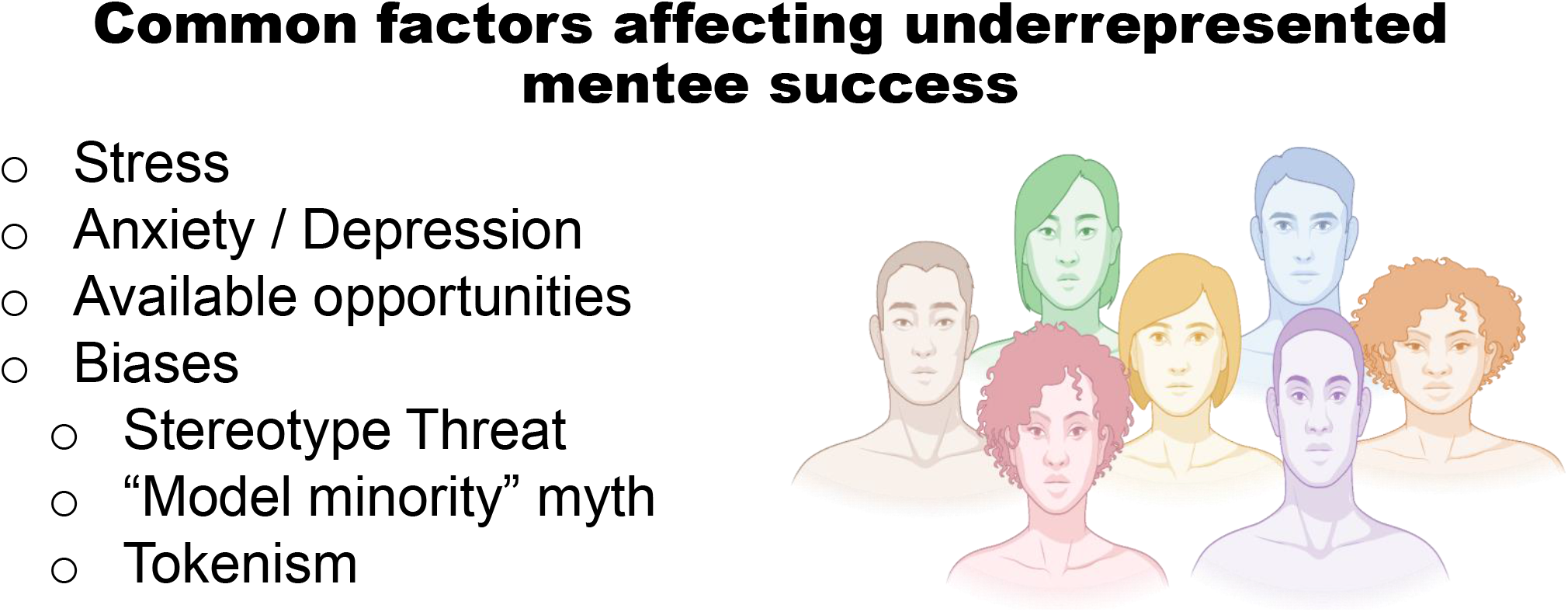
Graphic summarizing common factors affecting the success of underrepresented mentees in STEM, which can include stress, mental health issues like anxiety and depression, the availability of opportunities, and different types of biases that include stereotype threat, the myth of model minority, and tokenism.

**Figure 4.**
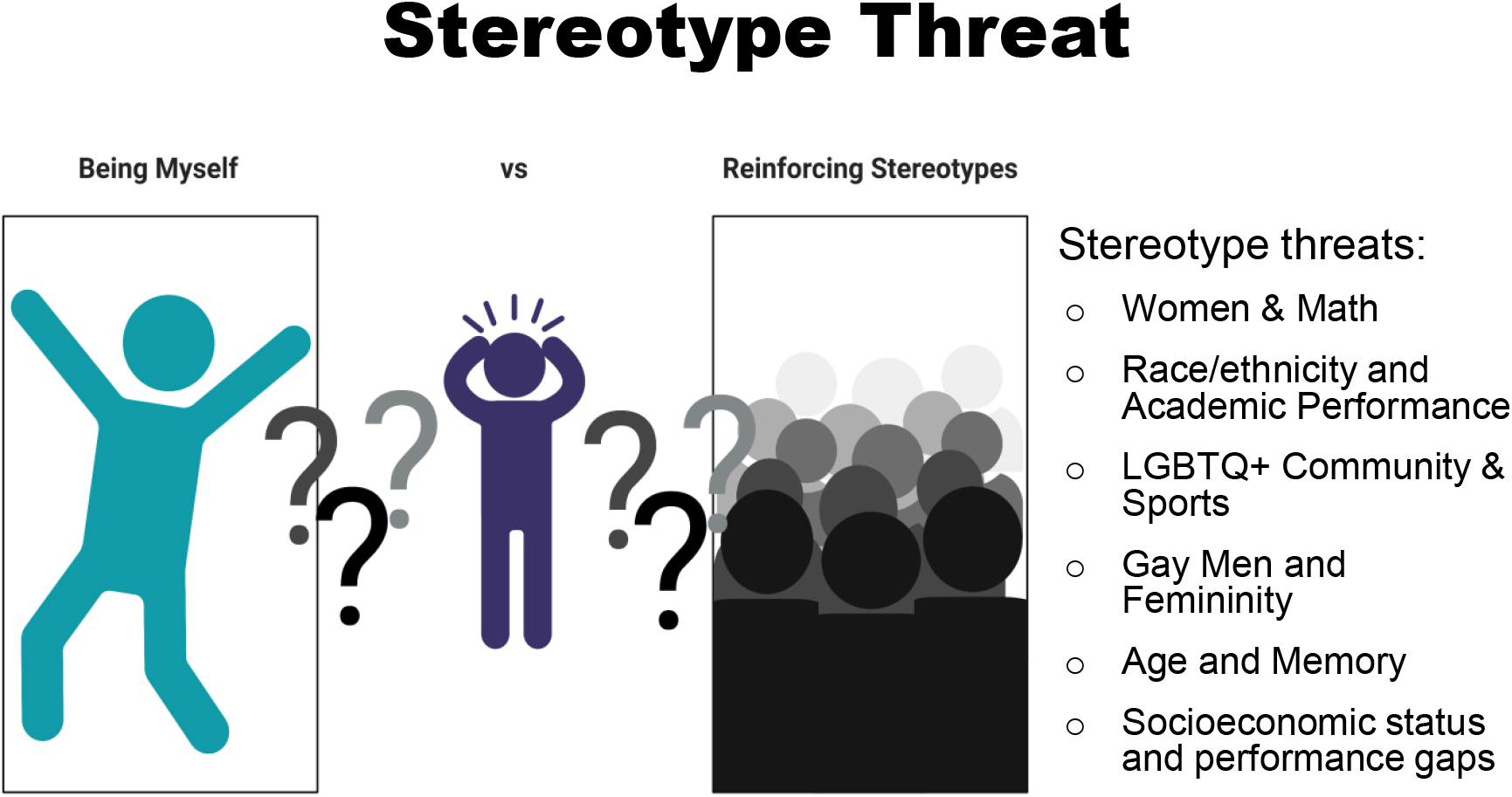
List of a common issues underlying stereotype threat that underrepresented individuals may have to deal with and overcome.

This seminar ultimately sought to explain the importance of mentors and how mentors should adapt for individual mentees to serve as an important resource. Mentors should be constantly shifting through getting out their message, boosting their mentees, and changing methods through the changing climates (Termini *et al*. 2021). For example, mentors should be able to adapt through electronic learning by e-mentoring through mediums such as video chatting, such as Skype or Zoom; or digital communication, including text, email, or GroupMe (McReynolds *et al*. 2020). For most of this workshop, these same points can be reversed – for example, for a mentee looking for a mentor, they may consider looking towards e-Mentoring.

## METHODS

We developed a one session workshop for 1.5 hours for participants regarding how to effectively produce quality work with multiple mentors. All of the data here was collected in response to a workshop given at Winston Salem State University to a sample (n) of 24 students from the general student population. This program may work with any collegiate students as the curriculum did not require any prerequisite knowledge.

Although we targeted first-year undergraduate students; this information may provide assistance for any stage of mentoring. The students who attended the workshop were given a pre/post-Likert scale-based survey to evaluate the effectiveness of the seminar on a wide range of topics including career and academic readiness.

### Workshop Overview

We designed this seminar to offer students career and academic direction in working with multiple mentors or principal investigators. We offered guidance on how to maintain healthy relationships with these mentors regardless of knowledge or prior interactions. Clips of prominent and diverse scientists or professionals with experience in mentoring offered various viewpoints to further the main points of this workshop. Case studies were utilized of other mentors from diverse groups to showcase the advantages of underrepresented mentors for mentoring. Discussions were designed around Socratic learning to encourage the use of critical thinking and connection to other topics such as the intrapersonal aspect of having, and working with, multiple mentors.

### Evaluating Feedback

Each student was administered a question pre/post-test to take independently (**Table 1**). Students were asked to rate (on a 10-point Likert-based scale, where 1 = *lowest* and 10 = *highest*) the effectiveness of the workshop. The pre/post-workshop questions asked students to rate their understanding, skills, and preparedness before and after the workshop. Understanding, skills, and preparedness were directly related to learning objectives from the workshop.

**Table 1.**
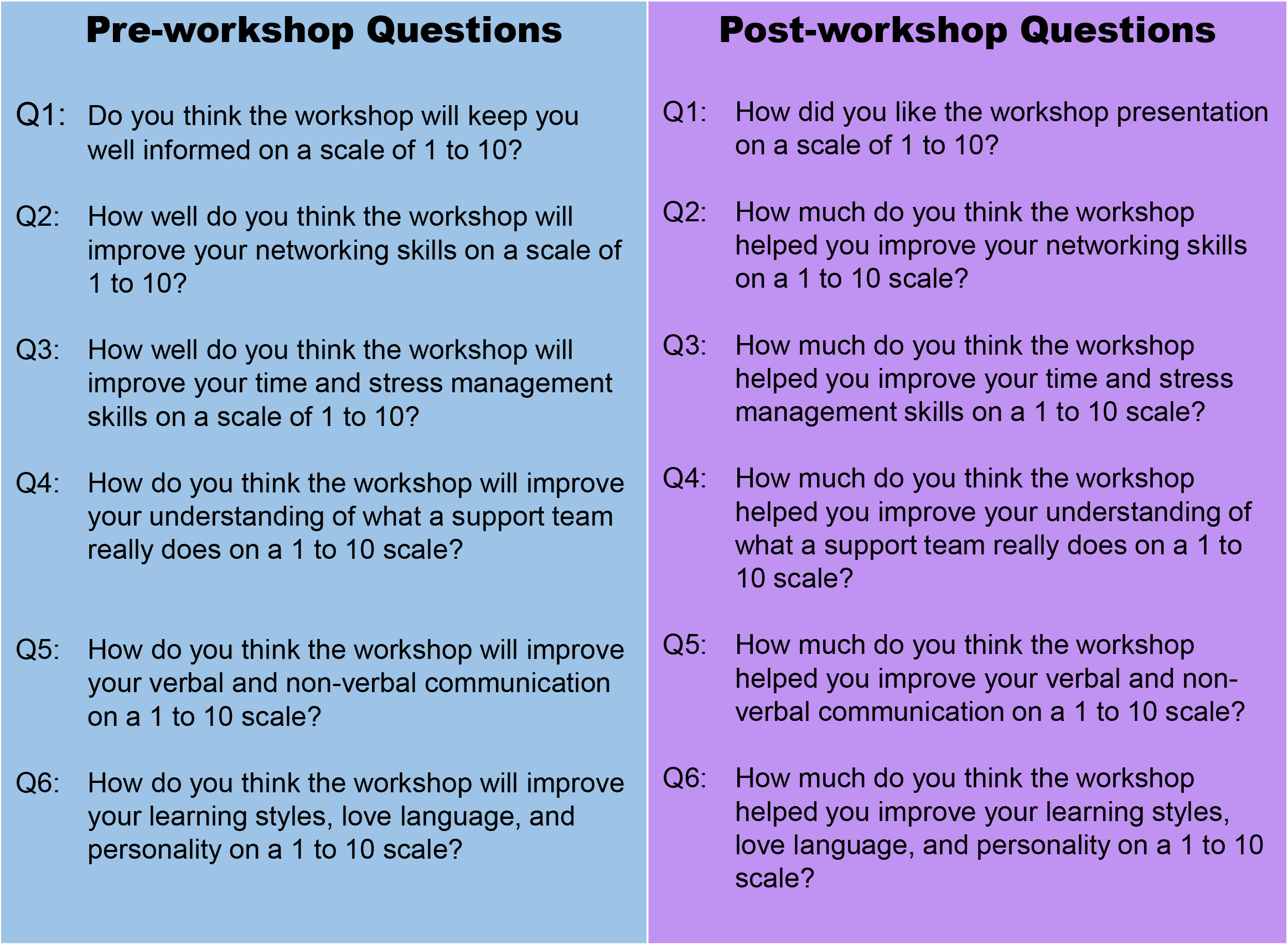
List of pre- and post-workshop questions administered to students.

### Statistical Analysis

The data for this study were collected via a pre/post-survey method. A Wilcoxon matched-pairs signed tank *t*-test analysis was use to determine the most effective to compare students’ answers before and after completion of the seminar. We summarized the data from the survey using box and whisker plots in which the centerline denotes the median, and error bars denote the standard error. Individual values are represented by circles. We considered differences statistically significant when *P* values were less than 0.05; NS (not significant). All analyses were performed using <u>GraphPad Prism 9.2.0</u>.

## RESULTS

Participating students were asked to complete a pre/post self-evaluation survey (*n* = 24, 100% response rate). All students were currently enrolled in the Historically Black College/University (HBCU) Winston Salem State University. Each evaluation question was designed in line with the objectives outlined for the workshop. Students initially did not see much value in the workshop, with an average score of 4.5 **(Figure 5 - Q1, pre-test)**. This intial low average may be due to lack of exposure to mentorship content or knowing what a mentor is beforehand; however, by the end of the workshop, the average score rose to 8.5 on a scale of 1–10, with 10 indicating the most favorable response **(Figure 5 – Q1, post-test)**. These results suggest that students found the workshop to be informative and have a better grasp of what is a mentor and what should they be looking for in a mentor. Prior to the workshop, students reported having little or no knowledge about networking skills with an average score of 4.0 out of 10 **(Figure 5 – Q2, pre-test)**. Despite this lack of knowledge on networking skills, participants found the workshop very informative and reported an average score of 9.4 out 10 at the end of the workshop **(Figure 5 – Q2, post-test)**, emphasizing the need for network training. On average participants reported a 3.9 score in whether they believed the workshop would improve their knowledge on time management skills; however by they end of the workshop their score rose to a 8.5 score **(Figure 5 – Q3)**, indicating a gain of knowledge on time and stress management. Paricipants also reported an averge score of 4.1 on whether or not the workshop would improve their understanding the role of support teams; however, by the end of the workshop participatants reported an average of 9.1, supporting the effectiveness of the workshop on learning about support teams **(Figure 5 – Q4)**. Participants felt, after engaging in the seminar, it helped them improve their non-verbal and verbal communication skills, **(Figure 5 – Q5)**. Intially participants seemed to have very little expectation on how the workshop would improve their learning styles, love language and personality with an average score of 2.9 and after attending the workshop students reported an average score of 9.2 score **(Figure 5 – Q6)**.

**Figure 5.**
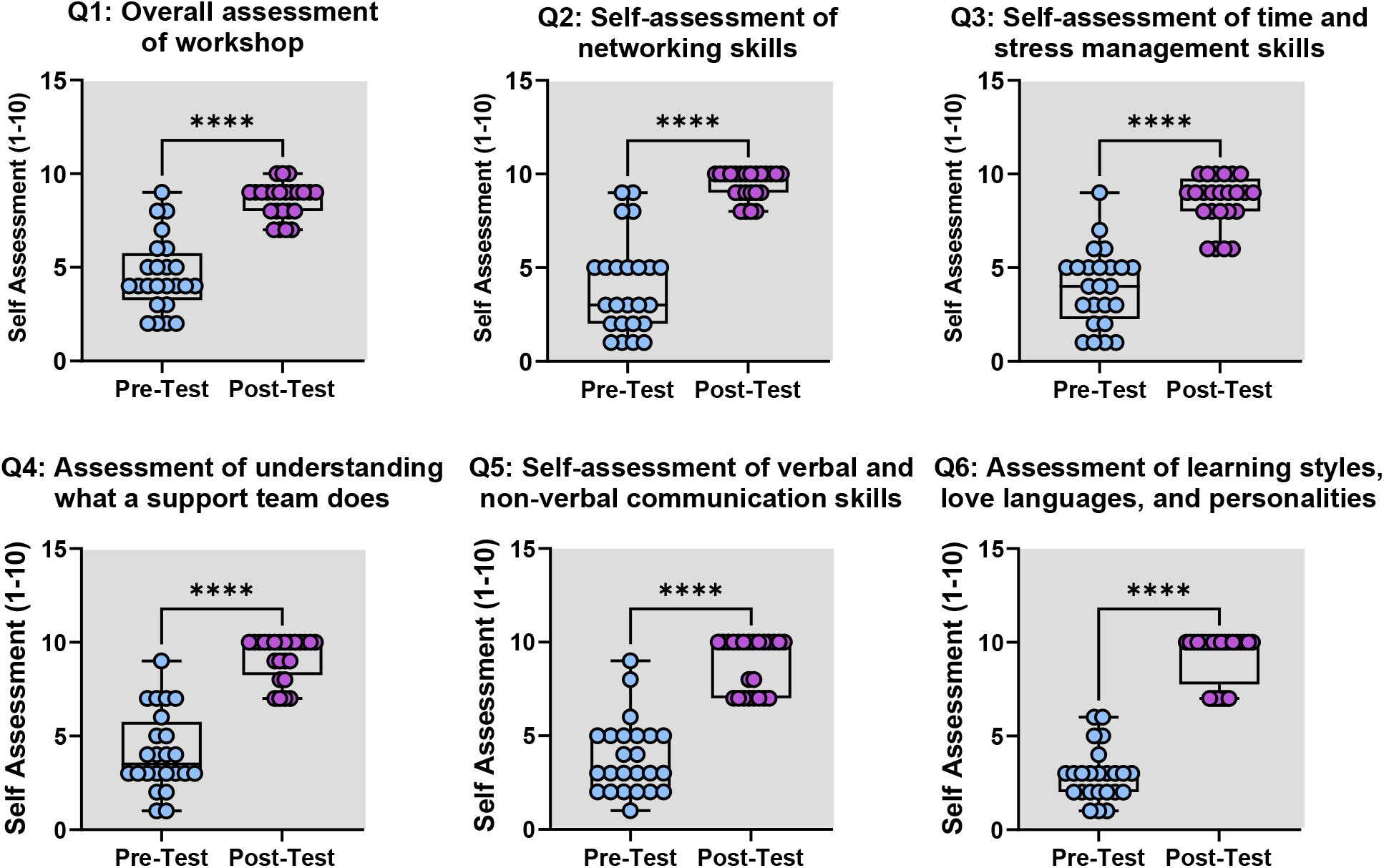
Qualitative summary of pre- and post-workshop questions administered to students.

## DISCUSSION

Having multiple mentors at each training stage is critical for student success. Many students are knowledgeable of the mentee-to-mentor relationship; however, students often lack a deeper understanding of how to best foster this relationship for ultimate academic achievement. Therefore, this workshop was developed to provide the necessary foundation for students to be better prepared for establishing future mentorships throughout graduate and professional school. For participants, the results of the workshop, “How to Handle More than One Mentor to Achieve Excellence,” indicated an overall enhancement of understanding mentorships as well as the importance of having multiple mentors. While there seemed to be little initial expectation that the seminar would produce anything beneficial, the participants ultimately felt like they gained knowledgeable information on a variety of topics related to the mentee-to-mentor relationship. Specifically, the pre/post-test questionnaire indicated appreciation for the effectiveness of the presentation on figuring out personal learning styles as well as how mentees prefer to be mentored. Additionally, participant responses suggested a better understanding of the importance of developing the necessary skills for successful matriculation into and through graduate and professional school.

The students that were a part of this exercise program were UR students from Winston Salem State University, indicating a diversity element to the effectiveness of this workshop. UR students often need direct, individualized support as well as a visual representation of successful mentors from similar backgrounds that understand what they are going through in academic spaces (Hinton Jr *et al*. 2020). To this end, the facilitators of the workshop provided several examples of successful mentee-to-mentor relationships, mentoring teams, as well as peer-mentor relationships of minoritized individuals. According to the pre/post-test survey, these examples in the presentation led to an increased understanding of what a support team does in general and for their mentees to achieve academic success.

## Limitations and Considerations

This workshop provided an adaptable virtual program to diverse pre-graduate and pre-professional students focused on the importance of establishing healthy mentorship relationships. The workshop was designed to instill confidence in students from marginalized backgrounds for future academic success with developing multiple mentee-to-mentor relationships. The virtual format makes this workshop extremely accessible to upcoming trainees in the academic pipeline as well as their potential mentors. Additionally, this program could be used as a refresher course to mentors on how to best support minoritized students both culturally and academically. However, this program had several limitations, including limited discussion between facilitators of the workshop and survey participants due to the pre-recorded virtual format, challenges with the type of evaluation and specific questions asked in the pre/post-survey, and potential for logistical difficulties.

The students assessed had a wide range of knowledge related to the topics discussed in the “How to Handle More than One Mentor to Achieve Excellence” workshop. As noted in the survey, some students had a stronger grasp of the issues presented in the workshops and therefore may have learned less from them. Overall, they felt as if the presentation would not be beneficial to them, but ultimately enjoyed the presentation. For a more thorough examination of the students’ assessment of the effectiveness of the workshop, survey questions could have been more specific to the topics discussed within the presentation. We did not conduct a focused pre-assessment of students to determine baseline levels of specific knowledge related to the topics discussed within the workshop. Inevitably, time spent on topics of most importance to the target audience may have been rushed when extra time to discuss was needed. Going forward, it will be critical to structure a more focused pre-assessment session before the workshop to make sure student knowledge of the intended topics are as equal as possible.

Future implementation of this workshop initiative can be adapted for individual institutions based on the needs of the intended audience, taking class size and demographics into account, as this may differ from the context of the original workshop. To allow for deeper engagement with the content of the workshop, adaptation of this workshop for future implementation could be expanded on the basis of the topics covered. Additionally, more specified questions could be asked in the pre/post-test survey which would help with overall understanding of the impact of the workshop on student learning and growth related to the topics covered.

Furthermore, this workshop could be adapted into a virtual format. The COVID-19 pandemic dramatically changed the way in which undergraduate and graduate students were educated and trained (Carr *et al*. 2021; Termini *et al*. 2021). The development and implementation of a virtual workshop to address issues related to students making use of multiple mentors, gaining networking experience and opportunities, and thoroughly understanding the importance of cultivating a healthy mentee-to-mentor relationship comes with its own challenges. However, adapting this existing workshop into a virtual format, while potentially limiting the opportunity for discussion between facilitators of the workshop and survey participants, is helpful to expanding the reach of the seminar. Especially critical to this, is the fact that many students, especially UR minorities, have faced increased academic difficulties due to COVID-19, highlighting the importance of mentoring (Gabster *et al*. 2020; Carr *et al*. 2021; Termini *et al*. 2021) Ultimately, we hope that this webinar will serve as a foundational body of work for mentors that may be beginning mentor/mentee relationships with underrepresented minoritized trainees in graduate or professional school.

## ACKNOWLEDGEMENTS

We would like to thank the students who participated in our workshop.

## Ethics Declaration

## Ethics Approval and consent to participate

## Consent for publication

