## Supplemental file for "The Importance of Mentors and How to Handle More Than One Mentor"

### How to Handle More than One Mentor to Achieve Excellence

### Educational Objectives

By the end of this session, participants will be able to

- Describe the role of mentors in developing the next generation of trainees
- Describe current research on mentorship among underrepresented populations.
- Apply skills on effective communication needed in the development of successful mentorship relationships.

### What is a Mentor ?

Consider inserting images and/or word art that symbolize good mentorship

Consider inserting images and/or word art that symbolize good mentorship

### What is a Mentor ?

- A willing and trusted advisor
- One who counsels and provides emotional support
- Mid 18$^{th}$ century: via French and Latin from Greek Mentor, the name of the adviser of the young Telemachus in Homer’s Odyssey

### What is a **Mentor** ?

First thing to do is find out who you are and what you want in life!

- Intentional
- Desire to Achieve
- Ability to Question
- Resolve
- Perseverance
- Access to great **verbal** and **non-verbal** communication
- To unlock your potential

Consider inserting a personal image of you and a mentor.

Insert inspirational quote here.

### Roles of a Mentor?

Consider inserting a word cloud image that encompasses all the roles of a mentor.

### Roles of a Mentor

- Advisor
- Coach
- Listener
- Encourager/Motivator
- Role Model
- Emotional Support

- Each has in common- Research Mentor

Consider inserting an image of someone being mentored/coached.

Insert inspirational quote here.

### The Traditional Mentor

- An advisor is responsible for career development

- Psychological support

Consider inserting a personal image of you and your mentees.

Insert inspirational quote here.

### The Educator and the Mentor

Consider inserting a personal image of you and your mentees.

- Great teachers serve the whole student.
- Across all ages, languages, ethnicities, and subjects, teachers are some of the most widely skilled people around in order to be successful.
- No matter what the goals are, they can pretty much be summed into a single sentence: You want to help people.
- And there are many ways you can help someone as a teacher. To name a few, teachers aspire to educate, to inspire, to learn and to affect positive change.

### The Educator and the Mentor

- Supports the improvement of scientific knowledge
- Trains one in the art of technical skills
- Enhances a trainees ability to critically and analytically think
- Identification of creative projects

Consider inserting a personal image of you and your mentees.

Insert inspirational quote here.

### The Supervisor

- Maintain performance
- Rectify behavior or encourage positive behavior
- Increase productivity

Consider inserting a personal image of you and your mentees.

Insert inspirational quote here.

### Coach

- Someone who is a life coach
- Develops skills that will help a person achieve life and career goals
- These skills vary depending on the type of mentor-to-mentee relationship

Consider inserting images of mentors with mentees.

Consider inserting images of mentors with mentees.

Consider inserting images of mentors with mentees.

### Sponsor

- One who provides funds or support for activities
- A career champion
- A sponsorship serves the mentor and the mentee

Consider inserting images of mentors with mentees.

### Sponsor

- Sponsorship, then, is a symbiosis rooted in action that furthers both sides' aspirations.

Consider inserting images of mentors with mentees.

### Mentoring/Role Models

- *Although "mentored residents were nearly twice as likely to describe excellent career preparation…under-represented minority residents were less likely to establish a mentoring relationship than their peers".*

- *Mentoring plays an important role in the perceived outcome on career readiness, making the difficulty underrepresented students face in developing a relationship with a mentor a serious .*

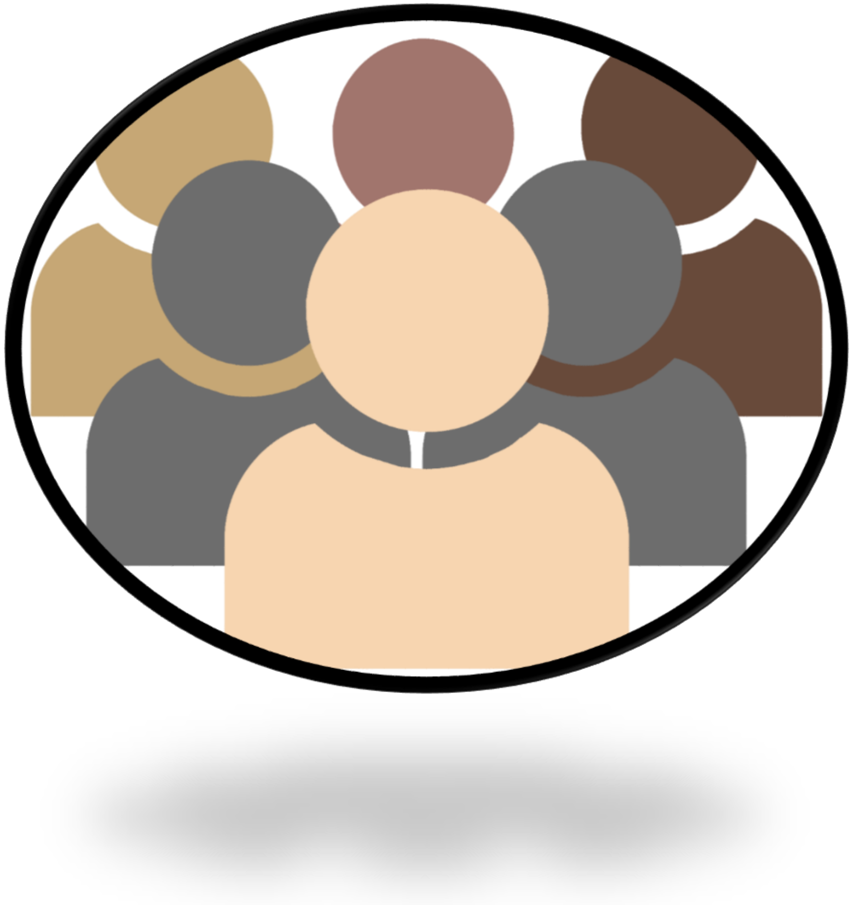

*Nivet MA. Minorities in academic medicine: Review of the literature. J Vasc Surg. Apr 2010;51(4 Suppl):53S-58S.*

### Mentoring/ Role Models

- *Mentoring programs should be tailored to their respective setting and environment–one-size fits all is not appropriate.*

- *"Mentoring must be* ***encouraged and rewarded*** *if underrepresented minorities are to achieve and maintain positions of influence and leadership in academia".*

*Lewellen-Williams, et. Al. The POD: A New Model for Mentoring Underrepresented Minority Faculty. Acad Med. 2006;81(3):275-279.*

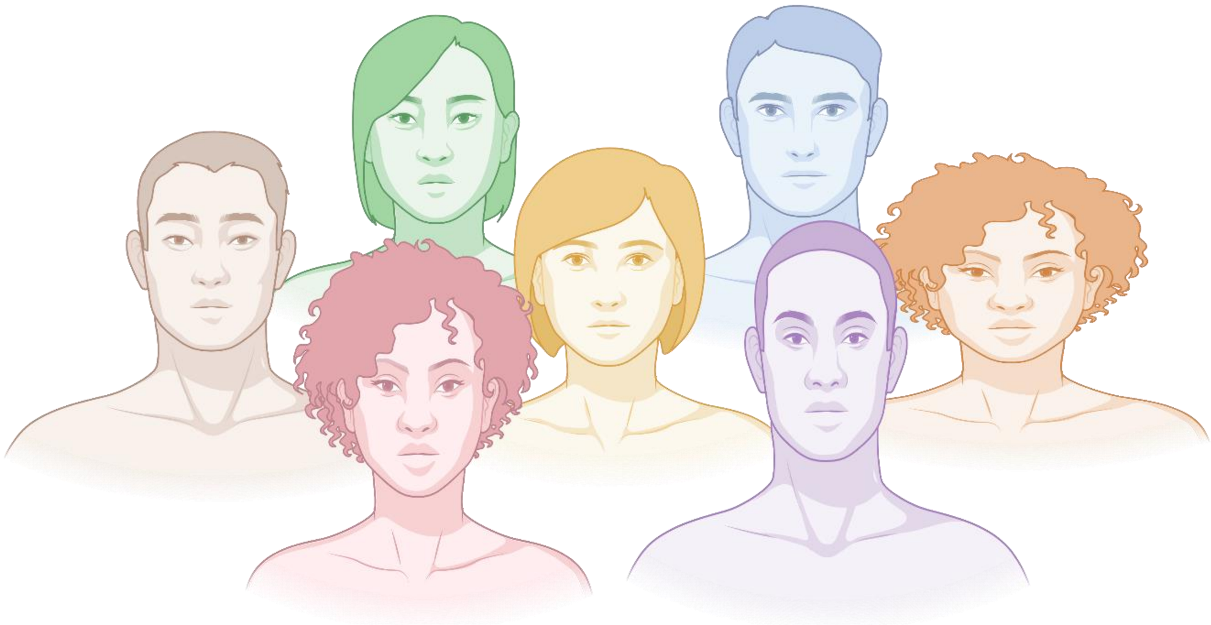

### Mentor-Mentee Concordance

- *Mentoring can play a significant role in addressing the lack of minority faculty and students at academic healthcare institutions as it provides "an avenue for interaction and camaraderie" amongst underrepresented students, faculty and staff*

- *While gender and ethnic similarities between the mentor and mentee are important factors, non-minority mentors "seem to understand and appreciate minority students' perspective and have volunteered to serve as mentors for these students"*

*Kosoko-Lasaki O, Sonnino RE, Voytko ML. Mentoring for women and underrepresented minority faculty and students: Experience at two institutions of higher education. J Natl Med Assoc.2006;98:1449–1459.*

Consider inserting images of mentors with mentees at a casual event.

### Factors that Influence Your Success

- High stakes situations

- Conflict around being a mentor and a manager
  - Stress
  - Anxiety
  - Deal with bullying in workplace
  - Lack of knowledge
  - Bias

Consider inserting fun lab images.

Consider inserting fun lab images.

Insert inspirational quote here.

### Important Things Mentors May Consider When Mentoring Underrepresented Groups

- Stereotype threat
- The affect  of stereotype threat
- Performance gaps

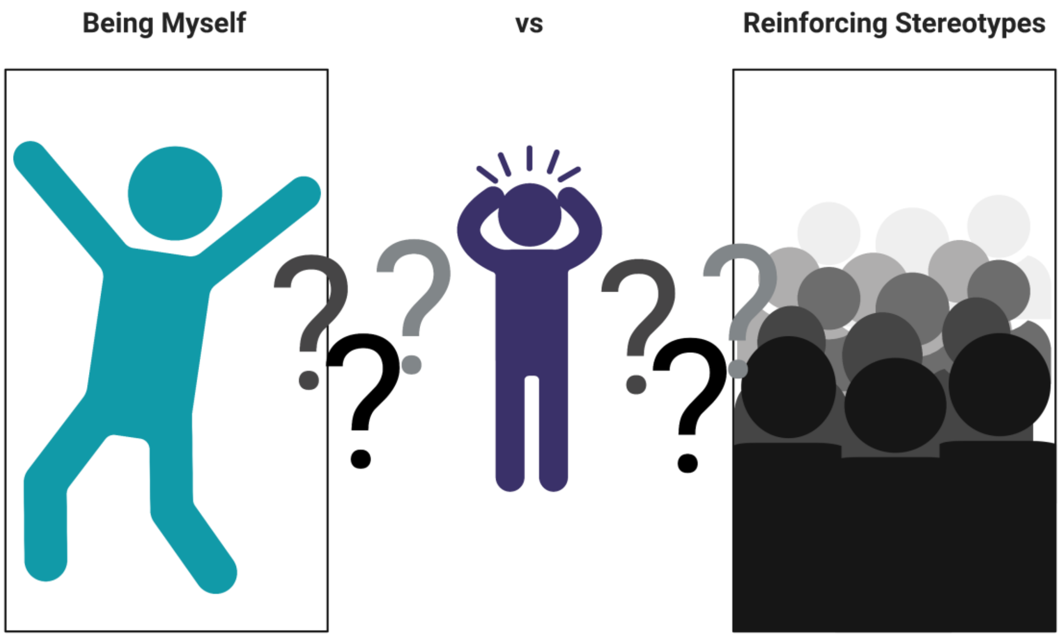


Insert inspirational quote here.

### Examples of Stereotype Threat

Stereotype threats:

- Women & Math
- African American & Academic Performance
- Latinx & Academic Performance
- Caucasians & Sports
- LGBTQ+ Community & Sports
- Gay Men and Femininity
- Age and Memory

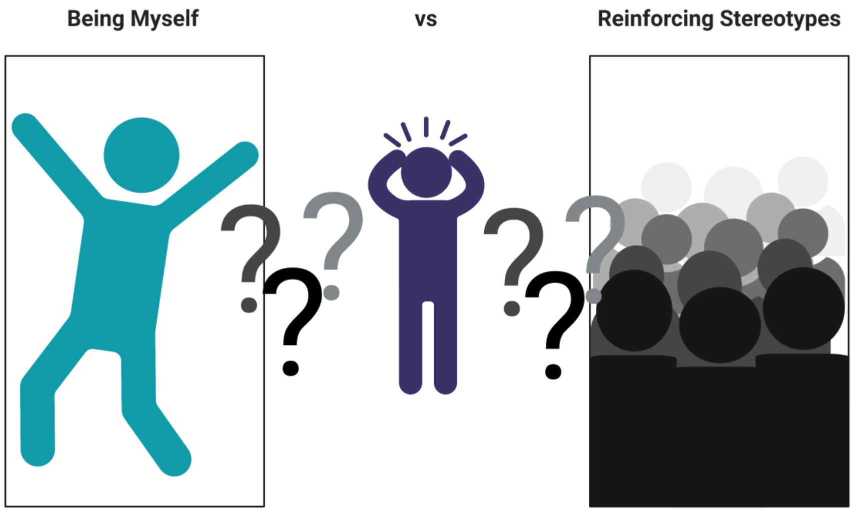


Social class, sexual orientation, and income, social class can all be stereotype threat issues.

Insert inspirational quote here.

### Examples of Stereotype Threat

#### Stereotype threat:

- The Claude Steele Study

*(Stereotype Threat and the Intellectual Test Performance of African Americans*

*Journal of Personality and Social Psychology, 1995, Vol. 69, No. 5, 797-811).*

- Race priming

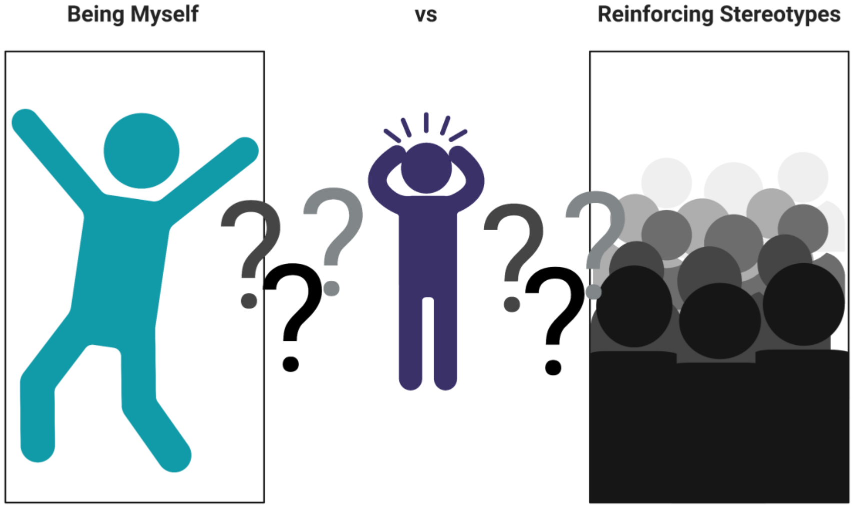


Insert inspirational quote here.

### Microaggressions

- Perceived incidents that were intended to cause subtle marginalization which can overtime have a negative impact on learning or performance.

  - Wow, your English is really good!
  - You should be good at this to a student form a certain group.
  - You didn't have to work twice as hard to get here.

Consider inserting images.

Consider inserting images.

Insert inspirational quote here.

### Macroaggressions

- **Reoccurring** incidents that are ***deliberate*** and immediate negative comments that impede learning or performance.

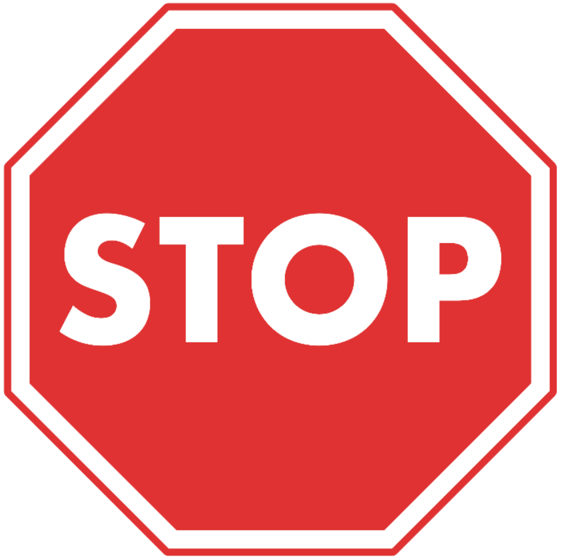


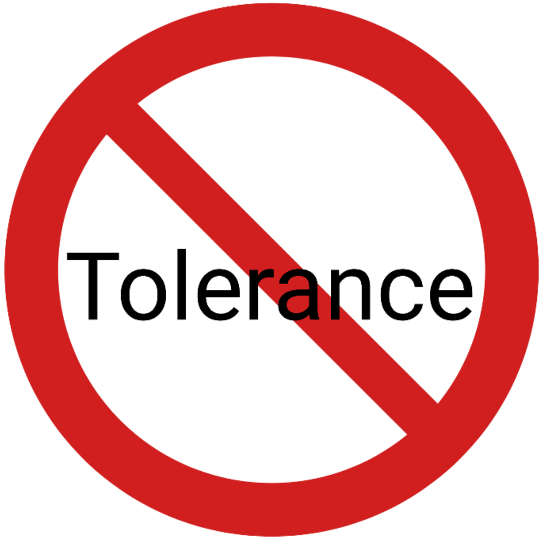

### Tokenism

- Defined as the model citizen
  - You ask the individual to express their ethnic or religious group's point of view for the class.
- Mentors should allow individuals to speak for themselves and not a group.
- This causes unnecessary harm

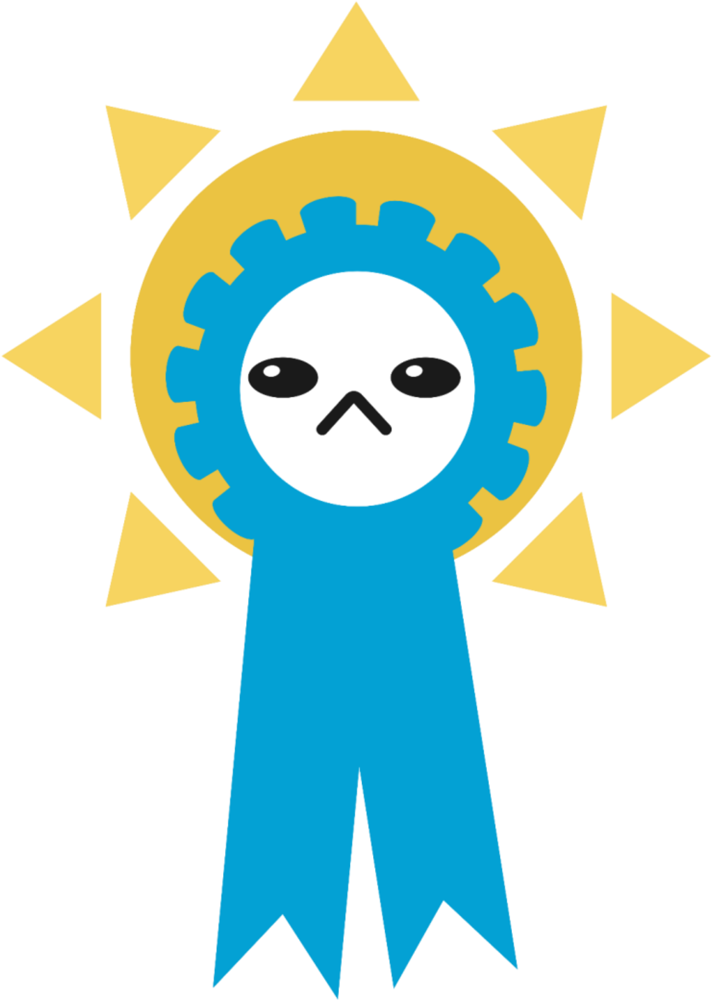

### Tokenism

#### Example:

In lab meeting,  a postdoctoral scholar  and a fellow are discussing some of the challenges they have experience with a collaborator. These collaborators are from Korea.

Both scholars have emailed the collaborator several times to ask them to send in their results, but received no response.

One of the lab mates turns to the only Korean person (who was not participating in the collaboration) and says, “Hey, Youngdo—you can probably tell me what’s going on, right?”

Insert inspirational quote here.

### Unconscious Biases

- ❖ Are prejudices in favor of or against one thing, person, or group compared with another, usually in a way that's considered to be unfair .

- ❖ Example, from a female Hispanic faculty perspective:

*"My office is near my community college faculty colleague's office. He is a male and doesn't have a doctoral degree. When people visit him and he is not there, they always assume:*

1) *That I am his secretary,  because I am a woman*
2) *That he has a doctorate, because he is a man"*

### What’s in it for me?

- A mentor can awaken your potential.
- A mentor can help establish strong networking skills.
- A mentor can help you establish trust with others .

Consider inserting images.

Consider inserting images.

Consider inserting images.

### What’s in it for me?

- Having the right network allows one to be able to carry out novel tasks and a myriad of diverse ideas.
- Being stagnant in one’s thinking leads to dangerous endings and does not advance one’s career. Innovation is what is needed to survive in a enigmatic economy.
- Key Question : How do I build new relationships?
  - First focus on what you can do: make a  Research gate, LinkedIn,  NIH National Research Mentoring Network, and a Professional Twitter.

### How to Build Strong Relationships

- Continued Key Question : How do I build new relationships?
- Find out how to be an engaging speaker by opening yourself up to listening to National Public Radio and Ted talks.
- Listening to Ted Talks will be able to find an example of what ever you are hoping to learn or deal with in the future?
- Example Ted Talk:

Consider inserting an example Ted Talk clip.

### How to Build Strong Relationships

- Conversation is also built on verbal and non-verbal communication skills.
- It is also important to  know when to use both.
- To explore verbal communication, we should set goals if we are trying to accomplish a transaction of some kind.

### Life Choices

- When you are working on achieving your goals one important thing is to be aware of other’s cultures.
- If you are aware of other’s cultures, you can also understand them better.
- Include an example where a lack of cultural awareness could lead to a misunderstanding.

### Life Choices

- Generally speaking, not being able to relate to others is the true causality.

- Usually, reaching goals means you need to acquire a team—which involves networking.
  - To be able to network, one must be willing to expand beyond their culture.

- Include an example of a common cultural custom/belief.

### IDP

- A mentor can help  boost confidence.

- Mentoring can improve your understanding of your goals and challenge individual growth and team goals made through an **IDP** or **SMART** goals.

### Personality

- Be willing to explore your personality!
- I would suggest learning who you are from the Myers Briggs Personality Test and the Big Five Personality Test.
- After focusing on your test, teach others how to love you. Take a Love Language test and in your networking and collaborations share who you are in your actions.

Consider inserting a graphic of the Meyers-Briggs personality types.

### Personality

* Next use your personality to mind your thought process.
* Building networks with others that are different than yourself.
* Diversity of Thought is important.
* Also, know your weaknesses of your personality.
* For example, if you start plans, but do not finish. Be willing to find out a way to influence yourself and be productive.

Consider inserting a graphic of the Meyers-Briggs personality types.

### Anxiety and Awfulizing

- How to be **respectful**, **spotting lies**, and passing **judgement**.
- A mentor can assist in building or developing a trademark/brand for yourself and mentor can teach you the dos and don'ts of **professional appearance**.
- A mentor can teach one to have mindfulness to decrease ones' stress and anxiety, improve time management, and **worry/awfulizing/catastrophizing.**

Consider including graphics of resources on these topics.

Consider including graphics of resources on these topics.

Consider including graphics of resources on these topics.

### Improvement

- A mentor improve your **body language**, **word choice**, and **diction**.
- A mentor can teach you **self-control** and how to assert **health boundaries**.
- A mentor can show you how to avoid anger during perceived **microaggressions** and **macroaggressions.**

Consider including graphics of resources on these topics.

Consider including graphics of resources on these topics.

Consider including graphics of resources on these topics.

### Boundaries

- Also, if you are unclear about boundaries, I suggest that you read books on it.
- An overview of how I establish boundaries:
  - Be truthful: No boundaries= little self esteem.
  - Decide your core values or concerns
  - What you will not compromise on for others
  - Decide the consequences ahead of time.
- Do not wavier from what you say.

Consider including a graphic of a resource on setting boundaries.

### Building a Brand/Trademark

Consider including a graphics of “branded” business cards, including your own if available. You may also consider references marketing for scientist resources.

### Factors that Enhance the Mentor-Mentee Relationship

- Strong EQ and IQ
- Focus
- Prioritize
- Capitalize
- Largest R.O.I.
- Stretch Goal to Stretch Experiences

Consider including a graphic you and your mentees.

### Body Language

* Body Language can be challenging to understand, but it is a component of non-verbal communication that can influence one's decision and actions.

* Therefore, it is important to not jump to conclusions with others too fast.

* Sit back and observe an individuals' behavior. The decorum of an individual will tell you a lot.

### Body Language

Consider including graphics that demonstrate how to read body language.

* Body Language can also deduce non-verbal communication about another individual, i.e. smiling

### Body Language

Consider including graphics that demonstrate how to read body language.

* Body Language can also deduce non-verbal communication about how someone focuses when delivering important information.

### Self-Control, Boundaries, Anger, and Sexual Harassment

* Being aware of one's self allows for a more productive environment.
* I suggest to find out if you have self-control or anger issues; try mediation and mindfulness training.
* Also be aware if you exhibit controlling behaviors.
* I also place harassment here because we all have to be aware of how we treat others and being realistic about proper training when interacting with others.

Consider including graphics workplace bullying and/or sexual harassment in the workplace. Or, include links to helpful resources related to self-control, anger, or controlling behaviors.

### Challenges to Models

- Institutional variability
- Resources
- Incentive models
- Specialties and subspecialties

### Mentor Search

- Institution
- Academic Society
- Regional Meetings
- Google
- Social Media
- Journals

Consider including a graphic of a mentor and mentee.

### Effective Meeting Checklist

- Location
- Clear aim or agenda set
- History &Physical
- Define Roles
- Personalization
- Goals and Expectations
- Setting dates
- Documentation
- Time & Tracking
- Summarize meeting

N A O'Dea, P de Chazal, D C Saltman, and M R Kidd. Running effective meetings: a primer for doctors. Postgrad Med J. Jul 2006; 82(969): 454–461. PMCID: PMC2563767

http://ctb.ku.edu/en/table-of-contents/leadership/group-facilitation/main

### Successful Mentoring Relationships

- **Preparation** is key to success
- Begin with the end in mind
- **Background** and **homework** prior to meeting
- Recurring Events
- Create time line
- **Consistency**
- Accessibility
- **Professionalism**
- BE IN THE KNOW!
- BE CURRENT!

### Mentoring Prep

Resources

- **Careers in Medicine**
- **Careers in Science**
- Specialty Website (AAS, AAOS, etc.)
- **SMART Goals**
- Current Schedule/ Curricula
- Resume/ CV
- **Transcript**
- Google
- IDP

### Technology & Mentoring

- E-Mentoring
  - Email
  - Text
  - Skype
  - Google Hangout

Consider inserting logos of top social media platforms, such as Twitter, FaceBook, LinkedIn, Slack, and GroupMe.

### Mentoring Challenges

- Reassess
- Refocus
- Recommit

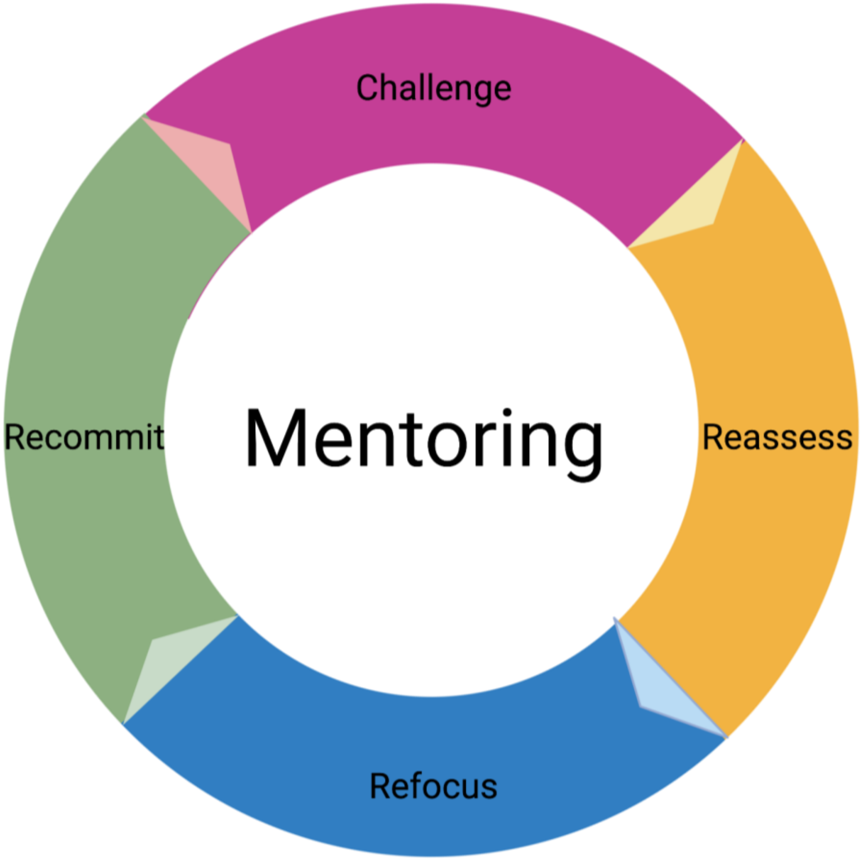

### Mentoring Cycle

- Practice of Mentoring
- Goal of outgrowing your mentor
- Investment of Mentor

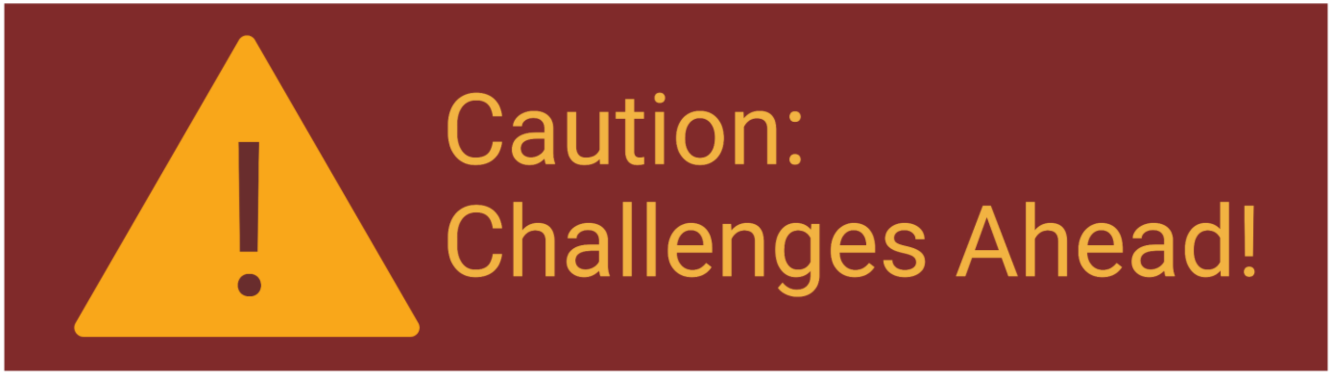
